# Persistence of the ground beetle (Coleoptera: Carabidae) microbiome to diet manipulation

**DOI:** 10.1101/2020.10.19.345207

**Authors:** Anita Silver, Sean Perez, Melanie Gee, Bethany Xu, Shreeya Garg, Kipling Will, Aman Gill

## Abstract

**Host-associated microbiomes can play important roles in the ecology and evolution of their insect hosts, but bacterial diversity in many insect groups remains poorly understood. Here we examine the relationship between host environment, host traits, and microbial diversity in three species in the ground beetle family (Coleoptera: Carabidae), a group of roughly 40,000 species that synthesize a wide diversity of defensive compounds**. This study found that the ground beetle microbiome is consistent across different host food sources. We used 16S amplicon sequencing to profile three species that are phylogenetically distantly related, trophically distinct, and whose defensive chemical secretions differ: *Anisodactylus similis* LeConte, 1851, *Pterostichus serripes* (LeConte, 1875), and *Brachinus elongatulus* Chaudoir, 1876. Wild-caught beetles were compared to individuals maintained in the lab for two weeks on carnivorous, herbivorous, or starvation diets. Soil environment but not diet had a significant effect on bacterial diversity and composition. The three carabid species have patterns of microbial diversity similar to those previously found in other insect hosts. Metagenomic samples from two highly active tissue types — guts, and pygidial gland secretory cells (which produce defensive compounds) — were processed and sequenced separately from those of the remaining body. The observed similarity of the pygidial gland secretory cell microbiome across hosts suggests the possibility that it may be a conserved community, possibly due to functional interactions related to defensive chemistry. These results provide a baseline for future studies of the role of microbes in the diversification of defensive chemical biosynthesis in carabids.

## INTRODUCTION

Insects are by far the most diverse group of animals (1) (2), and it is becoming clear that the success of several major insect groups is due in part to their resident microbiomes (2) (3). However, microbiomes and their possible symbiotic functions remain understudied in many major groups of insects, including Carabidae ground beetles. Carabidae consists of around 40,000 described species, making it one of the most species-rich animal families on earth (4). Moreover, the variety of defensive chemicals produced in the carabid pygidial gland system is an impressive example of evolutionary diversification (5). Secretory cells of the pygidial gland system produce such diverse classes of molecules as carboxylic acids, formic acid, quinones, hydrocarbons, and aromatics; chemical diversity exists even within some genera (5). Whether microbes play a functional role in carabid chemical diversity has not yet been studied.

Interactions between insects and their associated microbiomes can contribute to insect diversification (3). Microbiomes can benefit host insects in many ways, such as producing vitamin B (6), regulating host metabolism in response to stress (7), and contributing to host development (8). Notable examples of microbial symbionts supporting nutrient acquisition in insects include *Buchnera* bacteria producing essential amino acids allowing aphids to live on a nutrient-poor diet (9) and highly diverse termite gut microbes digesting cellulose for their wood-feeding hosts (10) (11). Unlike aphids and termites, carabids tend to be dietary generalists, and microbial species are also known to contribute to other host phenotypes, including nutrient acquisition and detoxification. In ants (12), *Harpalus pensylvanicus* (Degeer, 1774) (Carabidae) (13) and *Cephaloleia* (Coleoptera: Chrysomelidae) (14), microbial symbionts assist their hosts in metabolizing different food sources. It is known that bacterial symbionts enable several beetle species to thrive in chemically hostile environments. For example, the mountain pine beetle *Dendroctonus ponderosae* (Hopkins, 1902) (Coleoptera: Curculionidae) can inhabit pine trees because its microbes break down defensive terpenes produced by the trees (15). The microbiomes of *Nicrophorus vespilloides* Herbst, 1783 (Coleoptera: Silphidae) and other carrion beetles protect their hosts from toxins and speed up host digestion, making it easier for these beetles to feed on decaying carcasses (16) (17). Insects are well known to benefit from defensive and protective symbioses. *Lagria villosa* (F.) (Coleoptera: Tenebrionidae) beetles live in symbiosis with *Burkholderia gladioli* that protect their host’s eggs from pathogens by producing the antifungal compound lagriamide (18). *Paederus* (Coleoptera: Staphylinidae) beetles are well known for producing toxic hemolymph that causes severe dermatitis; the toxin, pederin, is produced by a *Pseudomonas*-like symbiont (19). The Asian citrus psyllid *Diaphorina citri* (Kuwafyama, 1908) (Hemiptera: Liviidae), an invasive pest in the U.S. that causes citrus disease, harbors endosymbiotic *Candidatus* Profftella armatura (Betaproteobacteria) that produce diaphorin, a toxin similar to pederin (20). Given the diversity of established insect-bacterial associations as well as the diversity of ground beetle defensive chemistry, it is worth examining the hypothesis that carabid microbiomes are associated with their defensive chemical production.

If bacteria indeed contribute to carabid host functions, we would expect a non-random microbiome composition corresponding to the functional role they play — in other words, microbial diversity should correspond to functional diversity. This is not necessarily straightforward, since insect microbiome composition can be explained by several factors such as host phylogeny (2) (21) (22) (23), dietary guild (23) or sampling locality (24). Although many insects have persistent host-associated communities, some do not, highlighting the potential for stochastic changes in microbial diversity; for example, some lepidoptera caterpillar microbiomes consist entirely of microbes ingested with leaves (25). If the carabid microbiome is similarly transient, it would be most closely resemble that of the community of the recent diet (12) (26) or other local environment (11), and not obviously correlate with other factors such as host phylogeny, chemistry, or tissue type. Host-associated microbiome composition can also be influenced by changes to host diet, as has previously been found in some Coleoptera and Lepidoptera species (14) (27). Provided that some of the variation in microbiome diversity and composition can be explained by factors outside of the transient aspects of the environment, however, it is possible that these microbes serve some function that benefits their host.

As insects have an open circulatory system that allows hemolymph to flow throughout the body, microbial communities are found in many insect tissues (28); but as in other animals, insects often have distinct microbial communities in different tissues (10) (12) (28). Some of this diversity may relate to the variety of conditions found within insect anatomy, including aerobic and anaerobic regions and extreme pH gradients (11) (28). Tissue-specific diversity could also be explained by a co-evolutionary relationship between hosts and symbiotic microbiota, in that hosts can harbor functionally useful bacteria in specialized tissues. For example, termites regulate unique microbiomes in each of several gut pouches (11), and many insect species maintain useful symbionts in specialized cells called bacteriocytes (28). The present study focuses on the pygidial gland secretory cells (hereafter just secretory cells) and the gut. We hypothesize that these tissues are most likely to host microbially-mediated functions because these organs are responsible for defensive chemical synthesis and digestion of food respectively, both known to involve bacterial symbionts in other insect taxa.

In this study, we used 16S metagenomic amplicon sequencing to quantify the bacterial diversity hosted by three carabid species under several dietary treatments. Each host species produces distinct primary defensive compounds: *Anisodactylus similis* produces formic acid, *Pterostichus serripes* carboxylic acids, and *Brachinus elongatulus* quinones (29)(Will & Attygalle, unpublished data). *Anisodactylus similus* has a distinct natural feeding preference from the other two species, so together these three species represent two different trophic types. *Brachinus elongatulus* and *P. serripes* are naturally generalist predator-scavengers, preferring animal matter but observed in nature and in the lab to eat a wide variety of sugar and protein rich plant and animal material; in contrast, *A. similis* is typically observed feeding on fallen fruits, seeds, and pollen (30) (Will unpbl.). In addition to sequencing wild-caught beetles preserved at the time of collection, we also subjected live beetles of each species to three specific dietary treatments, and dissected tissues of interest from each specimen.

This study examined how the transient factor of diet treatment, and more permanent factors including host species and tissue, contribute to the observed variation in carabid-associated microbiomes. We assume that if a given factor (e.g., food type or environment) does not impact the microbiome then the composition and diversity of the microbiome will appear random with respect to the state of that factor. On the other hand, if a factor is influential, we expect changes to that factor to significantly explain microbiome composition and diversity. Specifically, microbiome composition and diversity within treatments should be more similar than expected by random chance. We predicted that 1] If carabid beetles harbor non-transient host-associated microbes, then diet and local environmental factors would be insufficient to explain overall microbial variation across species. 2] Compared to diet treatment, host species would explain a greater share of microbial diversity — i.e. microbial communities would cluster more by host species than diet treatment. 3] A significant proportion of carabid microbial diversity would be explained by host tissue type. If microbial communities are random with respect to host tissue, that would undermine the hypothesis that microbes are involved in tissue-specific host functions like defensive chemical synthesis in secretory cells.

## METHODS

### Beetle husbandry and dissection

Twelve individuals each of *Anisodactylus similis* LeConte, 1851, *Pterostichus serripes* (LeConte, 1875), and *Brachinus elongatulus* Chaudoir, 1876 were collected (total 36 specimens). *Pterostichus serripes* and *A. similus* were collected from U.C. Berkeley’s Whitaker’s Forest, Tulare County, CA (36.7022°, −118.933°). *Brachinus elongatulus* were collected from national forest land in Madera Canyon, Santa Cruz County, AZ (31.72°, −110.88°). For each species, three wild-caught specimens were preserved in 95% ethanol immediately upon collection, and the remaining beetles were transported live to laboratory facilities on the U.C. Berkeley campus. For each species, in addition to wild-caught specimens, three diet treatments (banana, mealworm, and starvation) were tested in triplicate. Diet-treated beetles were kept in sterile containers with sterilized soil and water for 17 days in July, 2018. Banana-fed (Trader Joe’s, Dole Banana Ecuador) and mealworm-fed (Timberline, Vita-bugs Mini Mealworms 500 count) beetles were fed on the first day, and subsequently fed and watered every three days using heat-sterilized forceps and autoclaved water. All feeding portions were 0.04g (+/- 0.01g). Starved beetles received water, but no food. Banana and mealworm bacterial communities were sequenced as controls and were removed from the analysis after confirming samples were not contaminated. On the last day, beetles were quickly killed by placing them for one minute at −80°C in their plastic containers. All specimens, including wild-caught beetles, were dissected as described by McManus et al. (31). Each beetle was dissected into three groups of tissues: secretory cells, gut (including foregut, midgut, and hindgut), and the rest of the body minus the secretory cells and gut (subsequently referred to as ‘partial body’). Parasitic worms (Nematomorpha) found to be infecting one starved beetle and one mealworm-fed beetle were removed from those specimens and the worm tissues not included in downstream analysis.

### DNA extraction, PCR, and next generation sequencing

Tissues were incubated overnight in a 9:1 ratio of buffer ATL and proteinase K (Qiagen DNeasy Blood & Tissue Kit) at 55°C on a rocking tray. Lysate from overnight incubation was transferred to sterile 1.5ml O-ring tubes containing 0.25g (+/- 0.02g) of 0.1mm diameter zirconium beads and bead beat at 2000rpm for 3 minutes in a PowerLyzer to lyse bacterial cells. DNA was extracted from the lysed homogenate using Solid Phase Reversible Immobilization (SPRI) magnetic beads made following the method of Rohland (32): 100μL lysate was mixed with 180μL of well-mixed, room temperature SPRI beads, incubated for approximately 5 minutes on the bench, then transferred to a magnetic rack. After the SPRI beads pelleted, 200μL 80% ethanol was added. After 30 seconds the supernatant was removed, the ethanol wash was repeated a second time and the supernatant was removed again. Then, the tubes containing SPRI bead tubes were removed from the magnetic rack and allowed to air dry completely. DNA was eluted by adding 50μL TB solution (10mM Tris) directly onto the beads and incubating for 5 minutes, then returning samples to the magnetic rack to pellet the SPRI beads and retrieve the DNA-containing supernatant.

The V4 region of the 16S rRNA gene was PCR amplified in duplicate in 25μL reactions using GoTaq Green Master Mix (Promega), and the resulting PCR products were subsequently pooled. During the first round, previously described primers (33) 515FB_in (5’-ACA CTC TTT CCC TAC ACG ACG CTC TTC CGA TCT GTG YCA GCM GCC GCG GTA A-3’) and 806RB_in (5’-GTG ACT GGA GTT CAG ACG TGT GCT CTT CCG ATC TGG ACT ACH VGG GTW TCT AAT-3’), which were adapted to be complementary to the second round primers (34), were added to the ends of all 16S genes with the following conditions (BioRad thermocycler): initial denaturation at 94°C for 3 min, followed by 30 cycles of 94°C for 45 sec, 50°C for 1 min, 72°C for 1:30 min, and a final extension step of 72°C for 10 min. A second round of PCR was performed using unique combinations of barcoded forward (5’-AAT GAT ACG GCG ACC ACC GAG ATC TAC ACX XXX XXX XAC ACT CTT TCC CTA CAC GA-3’) and reverse (5’-CAA GCA GAA GAC GGC ATA CGA GAT XXX XXX XXG TGA CTG GAG TTC AGA CGT G-3’) primers (34) to create a dual-index amplicon library for Illumina sequencing (position of barcodes indicated by ‘X’ characters). The conditions for the second PCR reaction were: initial denaturation at 94°C for 3 min, followed by 10 cycles of 94°C for 45 sec, 50°C for 1 min, 72°C for 1:30 min, and a final extension step of 72°C for 10 min. All pooled duplicate PCR products were run on a 1% agarose gel for 30 min at 100V, and imaged under UV light to verify successful PCR. DNA concentration was quantified using a Qubit fluorometer, and equimolar amounts were pooled. The pooled library was purified (Qiagen Qiaquick PCR Purification Kit) and sent for Illumina MiSeq sequencing at the U.C. Berkeley Genomics Sequencing Laboratory.

### Analysis

Amplicon reads for the V4 region of 16S were de-multiplexed with deML (35) and processed using DADA2 (36), including quality filtering with maxEE=2. Reads were de-replicated into unique 16S amplicon sequence variants (ASVs, also referred to as phylotypes) using a read error model parameterized from the data. Paired-end reads were merged and mapped to ASVs to construct a sequence table. Chimeric sequences were removed. Taxonomic assignments for exact matches of ASVs and reference strains were made using the Ribosomal Database Project database (37). Sequence tables and taxonomic assignments were imported into R version 3.5 (38) for downstream analysis and combined into a single phyloseq (39) object for convenience. To account for variation in sequencing effort across samples, samples were scaled according to variance stabilized ASV abundances using DESeq2 (40) (41). ASV alignments made using DECIPHER (42) (43) were used to construct a neighbor-joining tree, and this tree was then used as the starting point for deriving a maximum likelihood tree from a generalized time-reversible model with gamma rate variation, implemented with the phangorn package in R (44). The tree was rooted using QsRutils (45). For comparative analysis between beetles, ASV data from all three tissues of each specimen were combined into an aggregate bacterial community. Alpha diversity measures were calculated using the packages phyloseq (39) and picante (46). Non-metric multidimensional scaling (NMDS) plots of beta diversity were created using phyloseq (39), and analysis of similarities (ANOSIM) tests were run using the package vegan (47). Bray-Curtis distances were calculated both for aggregate community data and for the original dataset. Venn diagrams of phylotypes present by diet were rendered by VennDiagram (48). To control for possible sequencing errors, only phylotypes occurring at least twice in the entire dataset were included in venn diagram analysis. Hierarchical clustering of communities was performed with the package ape (49). Secretory cells were tested for differential abundance of microbe phylotypes using an equivalent method to RNA-seq differential expression analysis, implemented using DESeq2 (40) (39).

### Ethics Statement

No permits were required for the described study, which complied with all relevant regulations.

## RESULTS

### Sequencing results

After quality filtering, the mean number of reads per sample was 19,868, and the median number of reads per sample was 17,532.

### Alpha diversity

There was a median of 95 and a mean of 98.6 ASVs present per sample.

### Diet

Phylogenetic diversity (PD) of aggregate communities was not associated significantly with diet treatment. When tissues were considered individually, only PD of the partial body varied significantly across diet treatments (Fig. 1). Richness results were similar to PD. Neither evenness nor Shannon diversity showed any significant effect of diet in either individual tissues or pooled microbiomes. Compared to wild-caught beetles, keeping the host in captivity subjected to any of the diet treatments had a minor effect on PD but not on other alpha diversity measures of the microbiome. Host diet did not correlate with community alpha diversity.

**Fig 1.**
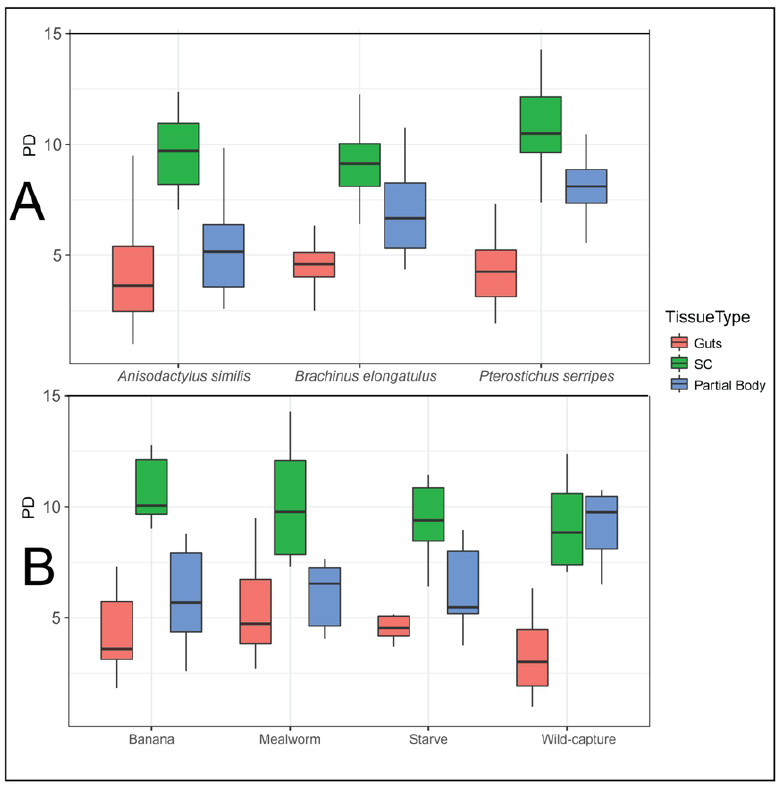
Boxplots of phylogenetic diversity (PD), with outliers depicted as points. (A) Plots grouped by host species. PD of partial bodies varied significantly by host species (chi-squared = 8.11, p = 0.017), but PD of all other tissues and of aggregate communities did not. (B) Plots grouped by diet treatment. PD of partial bodies varied significantly by diet treatment (chi-squared = 8.96, p = 0.030), but PD of all other tissues and of aggregate communities did not.

### Tissue

Tissue explained a large portion of the variance in PD of aggregate communities (Kruskal Wallis chi-squared = 55.5, p < 0.0001), so results were plotted separately for each tissue (Fig. 1). Secretory cell microbiomes had higher PD than gut microbiomes (Fig. 1). Richness, Shannon diversity, and evenness all varied significantly by tissue (p < 0.0001) as well.

### Species

Evidence of an effect of host species on microbial community diversity was relatively weak, and varied by tissue. Overall PD and evenness did not vary significantly by species. Richness (p = 0.043) and Shannon diversity (p = 0.041) varied only slightly significantly by species. Secretory cells had a very consistent alpha diversity level across species, only varying significantly by the measure of evenness (p = 0.030). Gut alpha diversity varied significantly across species by richness (p = 0.027), Shannon diversity (p < 1e-05), and evenness (p < 1e-05) but not PD. Differences in gut alpha diversity appear to be driven by the exceptionally low evenness in *A. similis* guts. The partial body microbiome had significantly different PD (Fig. 1) and richness (p = 0.0033), but no change in evenness, across host species. PD of partial bodies was highest in *P. serripes*.

### Community diversity distance analysis

Bray-Curtis distances for aggregate community data (Fig. 2a) and for the original dataset (Fig. 2b-d) reveal that community similarity is associated with several of the factors tested. Results of ANOSIM performed with Unifrac distances were consistent with results using Bray-Curtis distances reported below.

**Fig 2.**
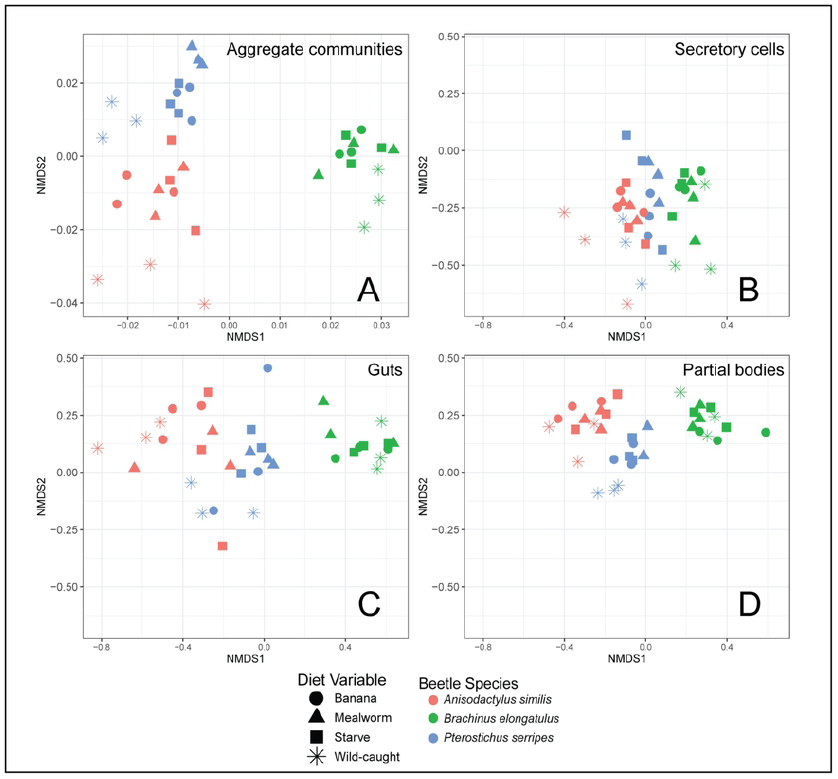
Bray-Curtis ordination of microbiome beta diversity using non-metric dimensional scaling. (A) Aggregate communities clustered significantly by species (ANOSIM R statistic = 0.92, p < 0.001) and tissue (R = 0.66, p < 0.001) only. (B) Secretory cell microbiomes clustered by species (R = 0.28, p < 0.001) and diet (R = 0.23, p < 0.001). (C) Gut microbiomes clustered clearly by species (R = 0.96, p < 0.001), and not by diet. (D) Partial body microbiomes are also clustered clearly by species (R = 0.95, p < 0.001), and not by diet.

### Diet

Aggregate communities did not cluster by diet (Fig. 2a). They did cluster by captive versus wild-caught beetles (ANOSIM R statistic = 0.3336, p < 0.001). The only tissue that clustered significantly by diet was secretory cells, but these clustered with a lower R statistic by diet (R = 0.23) than by species (R = 0.28). Clustering by diet was explained by significant differences between captive and wild-caught beetles. Phylotypes present in aggregate communities were compared across diet treatments (Fig. 3). A total of 1003 phylotypes were present across all diet conditions, 613 of which were present in wild-caught beetles. Of the phylotypes present in wild-caught beetles, 78% were present in at least one other diet condition. Just over a quarter of phylotypes were shared across all four diet conditions.

**Fig 3.**
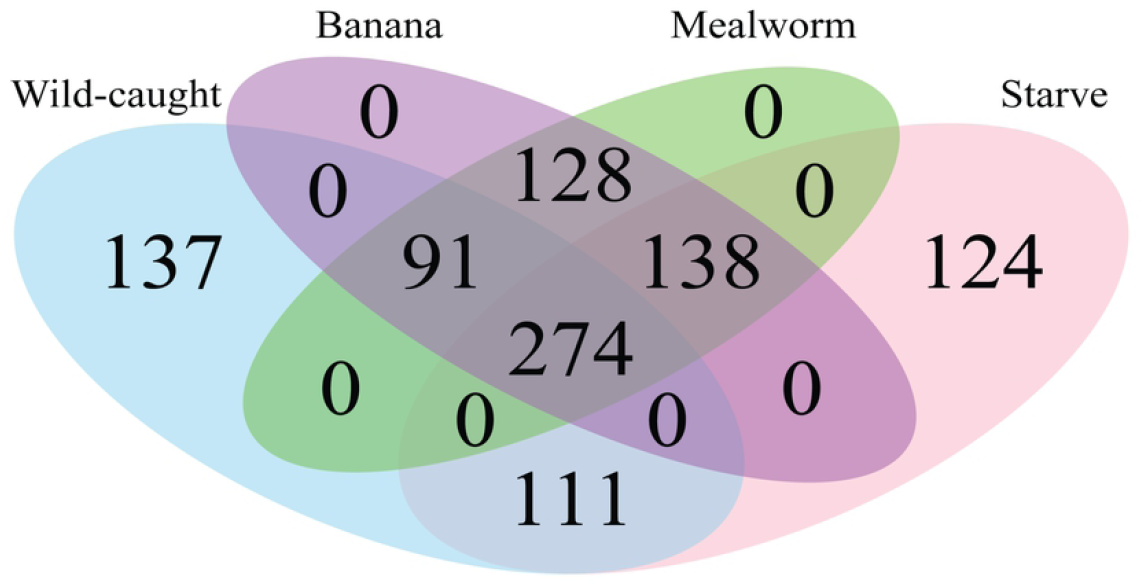
Venn diagram of phylotypes present in aggregate communities by diet treatment.

### Tissue

Microbial communities clustered clearly and significantly by host tissue.

### Specie

Aggregate communities clustered by host species, with the *B. elongatulus* microbiome being the most distinct (Fig. 2a). Individual tissues also clustered by species. The secretory cells had a much lower clustering statistic than the other tissues, indicating that microbial diversity in secretory cells is less differentiated than other tissues.

### Community Composition

The most abundant phyla across all samples were Proteobacteria (mean abundance 48.7%), Bacteroidetes (mean abundance 17.8%), Tenericutes, and Firmicutes (Fig. 4). Together, these four phyla comprised a mean of 94.6% of the bacteria in each sample. Communities in all beetle species and tissues had similar phylum-level compositions. Differences by host species arose more clearly at the level of bacterial genera, so community composition of each beetle species was plotted separately at this level (Fig. 4). Bacterial genera with median relative abundance across all samples of 1.5% or above were, in descending order of median relative abundance: *Acinetobacter, Spiroplasma, Yersinia, Flavobacterium, Pseudomonas, Enterobacter*, and *Enterococcus*.

**Fig 4.**
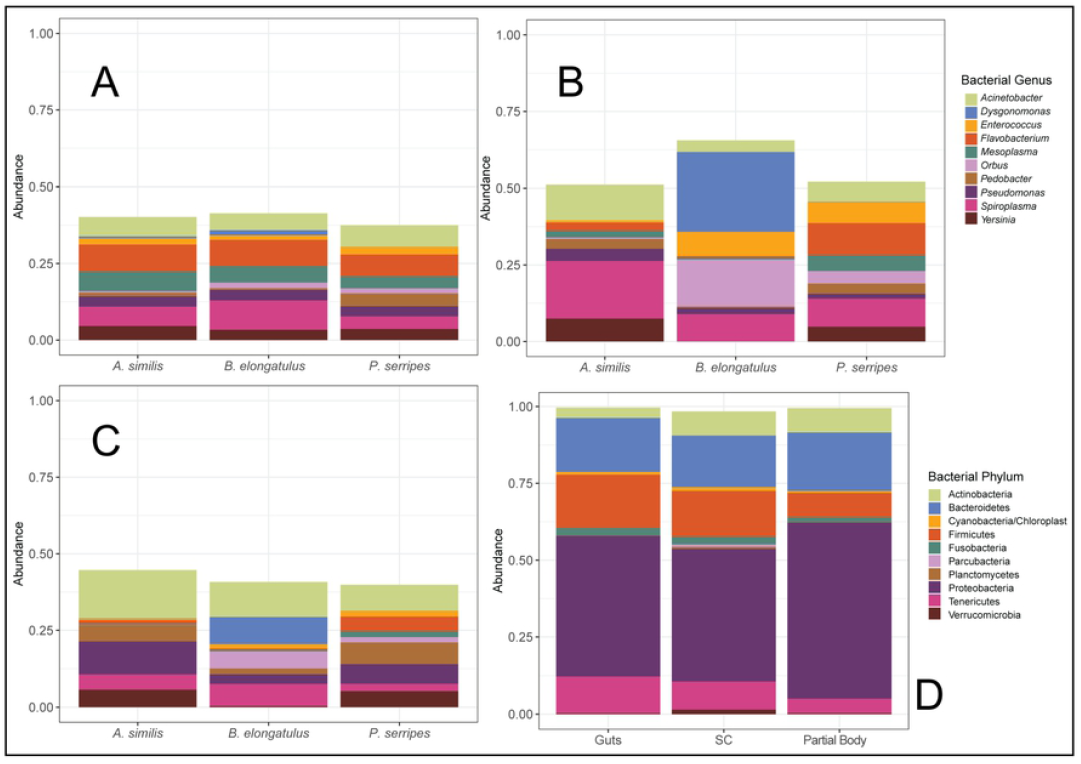
Relative abundances of prevalent bacterial taxa by host tissue and species. (A) Mean abundance in the secretory cells of the ten bacterial genera that were most abundant on average in all samples (n = 36, 12 per species). (B) Mean abundance of these bacterial genera in the guts (n = 36, 12 per species). (C) Mean abundance of these bacterial genera in the partial bodies (n = 36, 12 per species). (D) Mean relative abundance, across all host species, of the ten most abundant bacterial phyla (n = 108, 36 per tissue). Upper left legend applies to panels A-C and lower left to D.

### Diet

Community composition was not significantly different across diet treatments (Fig. 5).

**Fig 5.**
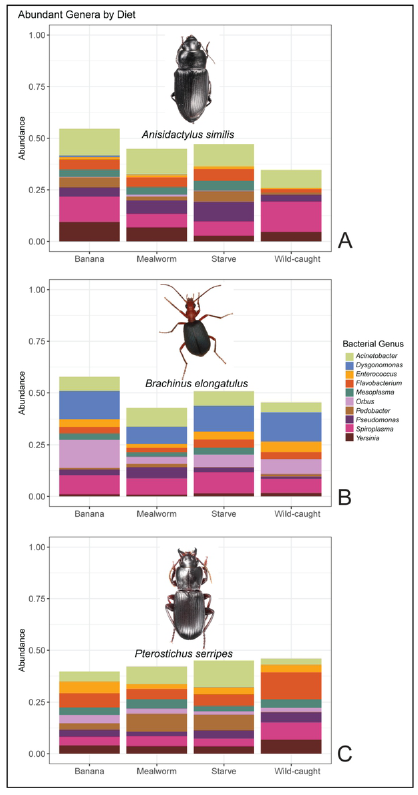
Relative abundances of prevalent bacterial taxa by host diet treatment. (A) Mean abundance in *A. similis* of the ten most abundant bacterial genera across all samples (n = 36), grouped by diet treatment (n = 9, each). (B) Mean abundance in *B. elongatulus* of these bacterial genera, grouped by diet treatment. (C) Mean abundance of these bacterial genera in *P. serripes*, grouped by diet treatment. Genera included are the same as in Fig 4. Photographs of beetles depict typical host morphology.

### Tissue

Differential abundance analysis of secretory cells versus all other tissues revealed that four phylotypes associated with two families were differentially expressed (p < 0.002). Two Flavobacterium phylotypes were more abundant in secretory cells than other tissues by factors of 10.22 and 16.23. Two Comamonadaceae phylotypes of unknown species were 9.38 and 9.92 fold more abundant in secretory cells. Secretory cell community composition is relatively conserved at the level of bacterial genera (Fig. 4). Compared to other tissues, gut microbiomes were more dominated by the ten most abundant bacterial genera; these ten genera composed over 50% of microbial abundance in all host species’ guts, over 60% of abundance in *B. elongatulus* guts, and less than 50% of abundance in other tissues’ microbial communities (Fig. 4).

### Species

Hierarchical clustering of community similarity showed that community differences corresponded with host species for all tissues. *Brachinus elongatulus* guts have more Firmicutes, and less Tenericutes and Actinobacteria, than the other two host species. Breaking down community composition to the genus level confirmed the status of *Brachinus* as the most distinct host species (Fig. 4).

## DISCUSSION

The present study assessed the degree to which diet and local environment shape microbiome composition and diversity in three carabid host species and among two active tissue types in each. Local environment was controlled for by comparing wild-caught and lab-reared beetles, and differences between these two conditions highlighted its importance. We hypothesized that if carabid beetle microbiomes play functional roles in chemical defense or other host traits, then a portion of microbial diversity should be non-transient; conversely, if carabid microbial diversity is entirely explained by diet and local environment, then it is unlikely microbes play a consequential role in the diversification of carabid defensive chemistry or other phenotypes. We found that patterns in microbiome composition and diversity are largely explained by the intrinsic factors of host species and tissue type. In contrast, shifts in host diet to carnivory, herbivory, or starvation had no significant effect on bacterial species diversity or composition. These findings demonstrate that carabid microbiomes are highly persistent to changes in host diet, paving the way for future efforts to decipher the ecological patterns and metabolic interactions that underlie non-transient host-associated microbial diversity in ground beetles. These results contribute to broader efforts to understand how the microbial diversity hosted by insects relates to insect evolution and ecology.

### Explanatory Power of Factors Tested

By subjecting beetles to differential dietary treatments in a controlled, sterile environment, this study quantified how transitory factors like host diet and soil environment predict variation in carabid microbial diversity, compared to intrinsic factors like tissue type and host taxonomy. The robustness of microbial community composition and diversity to diet changes, including to restricted herbivorous and carnivorous diets (Figs 1–3, 5), supports our first hypothesis of non-transient microbial communities. Controlling for local environment showed that soil is an important source of microbes for carabid beetles. As predicted in our second hypothesis, host species explains our results better than transient factors, both in terms of community composition (Fig. 4) and distance ordination. The finding that microbiome composition and diversity are associated most closely with tissue type strongly supports our third hypothesis. Tissue identity explains much of the variation between communities, in composition (Fig. 4) and distance ordination, and most of the variation in PD (Fig.1). Previous research has found that factors such as environmental filtering (3) and routes of microbe dispersal (22) can shape microbiome composition, and the especially strong association with tissue type could be related to these factors.

The random variation of microbial communities across diet treatments, together with the patterns in microbial community variation by host species and tissue type, indicates that carabid beetles possess non-transient, host-associated microbiomes. This is in accordance with our hypothesis that if carabid beetles harbor non-transient host-associated microbes, then diet and local environmental factors would not be sufficient to explain overall microbial variation across species. The finding that microbial communities are more similar within samples from the same host species supports our second hypothesis that microbes may be transmitted between individuals of the same species. The consistency of microbial communities within host species and tissue groups also raises the possibility that they co-evolved with their carabid hosts.

One transient factor that does explain a significant amount of the variation in our data is the impact of taking the host from the wild and keeping it under laboratory conditions on a common mix of sterile soil. Husbandry in a lab was previously found to have reduced microbiome diversity in carrion beetles (17) and lepidopteran species (27), indicating that a portion of these microbiomes are continuously acquired from the hosts’ surrounding environment. In the case of carabid beetles, soil may be an important source of microbial diversity. In our study, host captivity influences PD (Fig. 1) and the number of bacterial phylotypes present (i.e., richness) in individual microbiomes.

Factors other than those that were manipulated in this study could also exert influence on carabid microbiomes throughout the carabid life cycle. Our study does not address how juvenile carabids acquire microbes during their egg, larval, or pupal stages. Acquisition may occur by selective uptake from the environment as in leeches, hydra, and vibrio (11), or via direct transmission from a parent as in dung beetles (8). The relatively short time-frame of our study means it also cannot address the possible effects of long-term diet or environmental changes on carabid microbiomes, e.g., over the course of a season or year. Microbiome variability is explained by individuals’ long-term environment in drosophila (11) and houseflies (24). Finally, this study was not designed to disentangle the effects of host chemistry and host species identity, as each of the included species has different defensive chemical products.

### Comparison to other insect microbiomes

Given that related insect hosts sometimes share a subset of their microbiome, such as the conserved core microbiome of two dung beetles species (50), we anticipated that the three carabid species we tested might harbor distinct but overlapping microbial communities. Community distance ordination (Fig. 2), species- and genus-level composition, and similarity clustering showed that microbiomes were indeed distinct across *A. similis, P. serripes*, and *B. elongatulus*. The microbiomes of these three species also had many overlapping phylotypes.

A previous study of microbiome compositions showed that the resident gut microbiomes of two species of carabids, *Harpalus pensylvanicus* (Degeer, 1774) and *Anisodactylus sanctaecrucis* (Fabricius, 1798), had different composition and species richness from each other (51), and our results show a similar pattern of diversity across host species. The prevalence of the genus *Spiroplasma* in our results agrees with the findings of previous studies of carabid microbiomes (31) (51). *Dysgonomonas*, which has previously been found to be prevalent in *B. elongatulus* (31), was one of the most abundant genera, and also had higher relative abundance in *B. elongatulus* than in other beetle species, especially in the guts (Fig. 3). Another abundant genus in our study was *Enterococcus*, which is known to be associated with digestive tracts of several organisms including *B. elongatulus* (31). In *B. elongatulus* and *P. serripes*, we found *Enterococcus* to be more abundant in the guts than in the other tissues (Fig. 3).

The results of our small-scale study give additional preliminary evidence that carabid microbiomes are similar to other Coleoptera microbiomes. Several highly abundant bacterial genera in these samples were previously found in *Cephaloleia* (Chrysomelidae) beetles (14). The phylogenetic diversity of these carabid microbiomes is within the compass of previous studies in Coleoptera and in omnivorous insects (23). The phylogenetic diversity of the pygidial gland secretory cells, which are homologous structures found in carabids and other Adephaga, was unusually high. The lack of notable similarity in community composition between these results and previous studies of Coleoptera microbiomes confirmed past research that found that insect microbiome correspondence to phylogeny is not apparent when examined at deeper phylogenetic levels, typically corresponding to high-level taxonomy such as order (11) (21).

Comparing our results to past results across Insecta, we find that the five most abundant bacterial phyla in this study (Proteobacteria, Bacteroidetes, Actinobacteria, Tenericutes and Firmicutes) have previously been found to be the five most highly prevalent bacterial phyla in insects generally (2). Unlike what is known from gut samples from many other insect species (2), *Wolbachia* and *Rickettsia* are not among the most abundant bacterial genera in carabid guts. Insect gut microbiomes are more diverse across species than mammalian gut microbiomes (2), and past studies have shown that gut microbiomes group more loosely in beetles than in hymenopterans and termites (23). Comparison between the microbiomes from the carabid gut samples in our study and other insect gut microbiomes confirms this relatively high diversity. Gut phylotype richness in these samples was commensurate with the known range of richness among insect microbiomes (23), but higher than what has previously been found to be typical across several studies of insects (28).

### Possible functions of the microbes

Based on past studies of microbial symbiont functions, we hypothesized that microbes could play a role in carabid defensive chemical synthesis or nutrient metabolism. Specifically, that a significant proportion of carabid microbial diversity could be explained by host tissue type. The association found between host tissue type and patterns of microbiome composition and diversity tentatively support this hypothesis. This association, however, could also be a result of other functional relationships with the host, such as commensalism or parasitism, or simply an artifact of how the microbes are acquired.

There are no clear patterns in microbiome diversity specifically associated with guts (Fig. 4), so we cannot draw any conclusions about whether carabid gut microbes are involved in nutrient metabolism.

Microbial communities associated with secretory cells have much more similar composition (Fig. 2, Fig. 4) across host species than the composition associated with any other tissue. This similarity is noteworthy because secretory cells are a part of the pygidial gland system, an organ system which plays a conserved functional role across all host species. Possible explanations for this finding include coevolution or characteristics of the environment within secretory cells. This similarity is also consistent with the possibility that certain bacterial phylotypes play a symbiotic role in host chemical defense. To further investigate this last possibility, we checked for bacterial phylotypes that were differentially abundant in the secretory cells compared to other tissues. We found four such differentially expressed phylotypes: two from the genus *Flavobacterium* and two from an unidentified genus within the family Comamonadaceae. Although our study design limited our ability to draw definitive conclusions, it is distinctly possible that species that are especially abundant in the secretory cells could be involved in defensive chemical biosynthesis. Interestingly, some *Flavobacterium* species can produce quinones (52); *Flavobacterium* species are also known endosymbionts of giant scale insects (53).

### Future directions

This study found that patterns in microbial diversity and composition in carabid beetles are not random, and that the parameters that best explain them include host tissue and species. Transient changes in host diet have no significant effect on carabid microbiome diversity, although maintaining host beetles in sterile soil does have a modest but significant effect. Microbiome composition and diversity within the current, limited sample from across the carabid phylogeny appears to agree with previous findings regarding the microbiomes of Coleoptera and other insects.

Our results suggest that symbiosis may be a possibility, particularly in the secretory cells. Given the limitations of 16S amplicon data in assessing functional microbe-host interactions, future efforts to understand the nature of carabid microbiomes should consider shotgun metagenomic or other approaches that more directly quantify functional genes and metabolic pathways. Corroborating genetic data with experiments that confirm the metabolic activity of bacterial isolates from carabid tissues could also be useful. Additional future studies might use antibiotics to determine whether the presence of symbiotic microbes is essential for carabid host nutrition or defensive chemistry. Due to the great diversity among carabids, understanding the role of microbiomes in carabid hosts will be a key step toward understanding the diversity of possible host-microbiome interactions in insects and other animals.

## ACKNOWLEDGMENTS

We thank Wendy Moore and her lab group at the University of Arizona, Tucson, for providing the *Brachinus elongatulus* for the study.

## REFERENCES

1. Stork NE. How Many Species of Insects and Other Terrestrial Arthropods Are There on Earth? Annu Rev Entomol. 2018;63(1):31–45.

2. Yun J-H, Roh SW, Whon TW, Jung M-J, Kim M-S, Park D-S, et al. Insect Gut Bacterial Diversity Determined by Environmental Habitat, Diet, Developmental Stage, and Phylogeny of Host. Drake HL, editor. Appl Environ Microbiol. 2014 Sep 1;80(17):5254–64.

3. Vavre F, Kremer N. Microbial impacts on insect evolutionary diversification: from patterns to mechanisms. Curr Opin Insect Sci. 2014 Oct;4:29–34.

4. Lorenz W, 1950-. Systematic list of extant ground beetles of the world [Internet]. W. Lorenz; 2005 [cited 2019 Sep 27]. Available from: http://agris.fao.org/agris-search/search.do?recordID=US201300114567

5. Will KW, Attygalle AthulaB, Herath K. New defensive chemical data for ground beetles (Coleoptera: Carabidae): interpretations in a phylogenetic framework. Biol J Linn Soc. 2000 Nov;71(3):459–81.

6. Douglas AE. The microbial dimension in insect nutritional ecology. Funct Ecol. 2009 Feb;23(1):38–47.

7. Wong AC-N, Dobson AJ, Douglas AE. Gut microbiota dictates the metabolic response of Drosophila to diet. J Exp Biol. 2014 Jun 1;217(11):1894–901.

8. Schwab DB, Riggs HE, Newton ILG, Moczek AP. Developmental and Ecological Benefits of the Maternally Transmitted Microbiota in a Dung Beetle. Am Nat. 2016 Dec;188(6):679–92.

9. Douglas AE. Nutritional Interactions in Insect-Microbial Symbioses: Aphids and Their Symbiotic Bacteria *Buchnera*. Annu Rev Entomol. 1998 Jan;43(1):17–37.

10. Breznak JA. Intestinal Microbiota of Termites and other Xylophagous Insects. Annu Rev Microbiol. 1982 Oct;36(1):323–323.

11. Engel P, Moran NA. The gut microbiota of insects – diversity in structure and function. FEMS Microbiol Rev. 2013 Sep;37(5):699–735.

12. Russell JA, Moreau CS, Goldman-Huertas B, Fujiwara M, Lohman DJ, Pierce NE. Bacterial gut symbionts are tightly linked with the evolution of herbivory in ants. Proc Natl Acad Sci. 2009 Dec 15;106(50):21236–41.

13. Lundgren JG, Lehman RM. Bacterial Gut Symbionts Contribute to Seed Digestion in an Omnivorous Beetle. Van der Heijden M, editor. PLoS ONE. 2010 May 26;5(5):e10831.

14. Blankenchip CL, Michels DE, Braker HE, Goffredi SK. Diet breadth and exploitation of exotic plants shift the core microbiome of *Cephaloleia*, a group of tropical herbivorous beetles. PeerJ. 2018 May 17;6:e4793.

15. Adams AS, Aylward FO, Adams SM, Erbilgin N, Aukema BH, Currie CR, et al. Mountain Pine Beetles Colonizing Historical and Naïve Host Trees Are Associated with a Bacterial Community Highly Enriched in Genes Contributing to Terpene Metabolism. Appl Environ Microbiol. 2013 Jun 1;79(11):3468–75.

16. Vogel H, Shukla SP, Engl T, Weiss B, Fischer R, Steiger S, et al. The digestive and defensive basis of carcass utilization by the burying beetle and its microbiota. Nat Commun. 2017 Aug;8(1):15186.

17. Kaltenpoth M, Steiger S. Unearthing carrion beetles’ microbiome: characterization of bacterial and fungal hindgut communities across the Silphidae. Mol Ecol. 2014 Mar;23(6):1251–67.

18. Flórez LV, Scherlach K, Miller IJ, Rodrigues A, Kwan JC, Hertweck C, et al. An antifungal polyketide associated with horizontally acquired genes supports symbiont-mediated defense in Lagria villosa beetles. Nat Commun. 2018 Jun 26;9(1):1–10.

19. Kellner RLL. Molecular identification of an endosymbiotic bacterium associated with pederin biosynthesis in Paederus sabaeus (Coleoptera: Staphylinidae). Insect Biochem Mol Biol. 2002 Apr;32(4):389–95.

20. Nakabachi A, Ueoka R, Oshima K, Teta R, Mangoni A, Gurgui M, et al. Defensive Bacteriome Symbiont with a Drastically Reduced Genome. Curr Biol. 2013 Aug 5;23(15):1478–84.

21. Jones RT, Sanchez LG, Fierer N. A Cross-Taxon Analysis of Insect-Associated Bacterial Diversity. Gilbert JA, editor. PLoS ONE. 2013 Apr 16;8(4):e61218.

22. Brooks AW, Kohl KD, Brucker RM, van Opstal EJ, Bordenstein SR. Phylosymbiosis: Relationships and Functional Effects of Microbial Communities across Host Evolutionary History. Relman D, editor. PLOS Biol. 2016 Nov 18;14(11):e2000225.

23. Colman DR, Toolson EC, Takacs-Vesbach CD. Do diet and taxonomy influence insect gut bacterial communities? Mol Ecol. 2012 Oct;21(20):5124–37.

24. Bahrndorff S, de Jonge N, Skovgård H, Nielsen JL. Bacterial Communities Associated with Houseflies (Musca domestica L.) Sampled within and between Farms. Wilson BA, editor. PLOS ONE. 2017 Jan 12;12(1):e0169753.

25. Hammer TJ, Janzen DH, Hallwachs W, Jaffe SP, Fierer N. Caterpillars lack a resident gut microbiome. Proc Natl Acad Sci. 2017 Sep 5;114(36):9641–6.

26. Scully ED, Geib SM, Carlson JE, Tien M, McKenna D, Hoover K. Functional genomics and microbiome profiling of the Asian longhorned beetle (Anoplophora glabripennis) reveal insights into the digestive physiology and nutritional ecology of wood feeding beetles. BMC Genomics. 2014;15(1):1096.

27. Tang X, Freitak D, Vogel H, Ping L, Shao Y, Cordero EA, et al. Complexity and Variability of Gut Commensal Microbiota in Polyphagous Lepidopteran Larvae. Dale C, editor. PLoS ONE. 2012 Jul 17;7(7):e36978.

28. Douglas AE. Lessons from Studying Insect Symbioses. Cell Host Microbe. 2011 Oct;10(4):359–67.

29. Attygalle AB, Xu S, Moore W, McManus R, Gill A, Will K. Biosynthetic origin of benzoquinones in the explosive discharge of the bombardier beetle Brachinus elongatulus. Sci Nat. 2020 Aug 1;107(4):1–11.

30. Larochelle A. The food of carabid beetles (Coleoptera: Carabidae, including Cicindelinae). Fabreries. 1990;Supplément 5:1–132.

31. McManus R, Ravenscraft A, Moore W. Bacterial Associates of a Gregarious Riparian Beetle With Explosive Defensive Chemistry. Front Microbiol. 2018 Oct 5;9:2361.

32. Rohland N, Reich D. Cost-effective, high-throughput DNA sequencing libraries for multiplexed target capture. Genome Res. 2012 May 1;22(5):939–46.

33. Caporaso JG, Lauber CL, Walters WA, Berg-Lyons D, Huntley J, Fierer N, et al. Ultra-high-throughput microbial community analysis on the Illumina HiSeq and MiSeq platforms. ISME J. 2012 Aug;6(8):1621–4.

34. Lange V, Böhme I, Hofmann J, Lang K, Sauter J, Schöne B, et al. Cost-efficient high-throughput HLA typing by MiSeq amplicon sequencing. BMC Genomics. 2014 Jan 24;15:63.

35. Renaud G. grenaud/deML [Internet]. 2019 [cited 2019 Sep 25]. Available from: https://github.com/grenaud/deML

36. Callahan BJ, McMurdie PJ, Rosen MJ, Han AW, Johnson AJA, Holmes SP. DADA2: High-resolution sample inference from Illumina amplicon data. Nat Methods. 2016 Jul;13(7):581–3.

37. Cole JR, Wang Q, Fish JA, Chai B, McGarrell DM, Sun Y, et al. Ribosomal Database Project: data and tools for high throughput rRNA analysis. Nucleic Acids Res. 2014 Jan 1;42(Database issue):D633–42.

38. R Core Team. R: A language and environment for statistical computing [Internet]. Vienna, Austria: R Foundation for Statistical Computing; 2018 [cited 2019 Sep 25]. Available from: https://www.R-project.org/

39. McMurdie PJ, Holmes S. phyloseq: An R Package for Reproducible Interactive Analysis and Graphics of Microbiome Census Data. PLOS ONE. 2013 Apr 22;8(4):e61217.

40. Love MI, Huber W, Anders S. Moderated estimation of fold change and dispersion for RNA-seq data with DESeq2. Genome Biol. 2014;15(12):550.

41. McMurdie PJ, Holmes S. Waste Not, Want Not: Why Rarefying Microbiome Data Is Inadmissible. PLOS Comput Biol. 2014 Apr 3;10(4):e1003531.

42. Wright ES. DECIPHER: harnessing local sequence context to improve protein multiple sequence alignment. BMC Bioinformatics. 2015 Oct 6;16(1):322.

43. Wright E S. Using DECIPHER v2.0 to Analyze Big Biological Sequence Data in R. R J. 2016;8(1):352.

44. Schliep KP. phangorn: phylogenetic analysis in R. Bioinformatics. 2011 Feb 15;27:592–3.

45. Quensen J. QsRutils: R Functions Useful for Community Ecology [Internet]. 2019 [cited 2019 Sep 25]. Available from: https://github.com/jfq3/QsRutils

46. Kembel SW, Cowan PD, Helmus MR, Cornwell WK, Morlon H, Ackerly DD, et al. Picante: R tools for integrating phylogenies and ecology. Bioinformatics. 2010 Jun 1;26(11):1463–4.

47. Oksanen J, Blanchet FG, Friendly M, Kindt R, Legendre P, McGlinn D, et al. vegan: Community Ecology Package [Internet]. 2019 [cited 2019 Sep 25]. Available from: https://CRAN.R-project.org/package=vegan

48. Chen H. VennDiagram: Generate High-Resolution Venn and Euler Plots [Internet]. 2018 [cited 2019 Sep 25]. Available from: https://CRAN.R-project.org/package=VennDiagram

49. Paradis E, Schliep K. ape 5.0: an environment for modern phylogenetics and evolutionary analyses in R. Bioinformatics. 2019 Feb 1;35(3):526–8.

50. Franzini PZN, Ramond J-B, Scholtz CH, Sole CL, Ronca S, Cowan DA. The Gut Microbiomes of Two Pachysoma MacLeay Desert Dung Beetle Species (Coleoptera: Scarabaeidae: Scarabaeinae) Feeding on Different Diets. Forster RJ, editor. PLOS ONE. 2016 Aug 17;11(8):e0161118.

51. Lundgren JG, Lehman RM, Chee-sanford J. Bacterial Communities within Digestive Tracts of Ground Beetles (Coleoptera: Carabidae). Ann Entomol Soc Am. 2007 Mar 1;100(2):275–82.

52. Collins MD, Ross HNM, Tindall BJ, Grant WD. Distribution of Isoprenoid Quinones in Halophilic Bacteria. J Appl Bacteriol. 1981 Jun;50(3):559–65.

53. Matsuura Y, Koga R, Nikoh N, Meng X-Y, Hanada S, Fukatsu T. Huge Symbiotic Organs in Giant Scale Insects of the Genus Drosicha (Coccoidea: Monophlebidae) Harbor Flavobacterial and Enterobacterial Endosymbionts. Zoolog Sci. 2009 Jul;26(7):448–56.

